# Neuronal memory for language processing

**DOI:** 10.1101/546325

**Authors:** Hartmut Fitz, Marvin Uhlmann, Dick van den Broek, Renato Duarte, Peter Hagoort, Karl Magnus Petersson

**Affiliations:** Donders Centre for Cognitive Neuroimaging, Radboud University Nijmegen, the Netherlands; Neurobiology of Language Department, Max Planck Institute for Psycholinguistics Nijmegen, the Netherlands; Institute of Neuroscience and Medicine (INM-6) and Institute for Advanced Simulation (IAS-6) and JARA-Institute Brain Structure-Function Relationships (INM-10), Jülich Research Centre, Germany; Centre for Biomedical Research, University of Algarve, Portugal

## Abstract

In language processing, an interpretation is computed incrementally within memory while utterances unfold in time. Here, we investigate the nature of this processing memory in a spiking network model of sentence comprehension. We show that the history dependence of neuronal responses endows circuits of biological neurons with adequate memory to assign semantic roles and resolve binding relations between words in a stream of language input. A neurobiological read-write memory is proposed where short-lived spiking activity encodes information into coupled dynamic variables that move at slower timescales. This state-dependent network does not rely on persistent activity, excitatory feedback, or synaptic plasticity for storage. Instead, information is maintained in adaptive neuronal conductances and can be accessed directly during comprehension without cued retrieval of previous input words. This work provides a step towards a computational neurobiology of language.

## Introduction

Language processing requires memory ranging from milliseconds to seconds and minutes, and virtually every psycholinguistic theory—from acquisition to adult processing and language disorders—implicates the structure of this system. Prominent theories of memory (Baddeley, 1986; Just and Carpenter, 1992; Lewis et al., 2006), however, have made little contact with the detailed neurobiological infrastructure that implements language. On the other hand, theories of short-term memory developed in neuroscience are often based on simple delayed response tasks (Goldman-Rakic, 1995; Fuster, 1997; Mongillo et al., 2008) while memory requirements for language processing differ substantially from this paradigm. During comprehension, linguistic units are integrated into a sentence-level interpretation. Cues to sentence meaning include lexical semantics, morphology and syntax, and these can occur anywhere in the input, and at variable distance from the location where they are being used. For instance, in *“the cheese is eaten by the mouse”*, noun animacy (*cheese, mouse*), verb identity (*eat*), inflectional morphemes (-*en*) and function words (*by*) jointly support an interpretation of *mouse* as the AGENT of the action. Integration of cues takes place in an incremental fashion, as soon as they become available to the processing machinery. Input is not just stored passively in a buffer and loaded back into the comprehension system when needed. Instead, the meaning of an utterance is constructed *within* online processing memory as it unfolds in time, suggesting that memory for language is actively computing (Petersson and Hagoort, 2012). Secondly, sentences have internal structure and the order of word input matters (e.g., *“cat chases dog”* versus *“dog chases cat”*). Thus, processing memory for language needs to be sensitive to precedence relations and it is currently unknown how this is achieved in neural circuits. For instance, simple attractor-based models are not capable of maintaining the order of sequential input because temporal information is lost (Ganguli et al., 2008). And third, comprehension is a rapid process (2–4 words per second in listening, 4–6 in reading; Rayner and Clifton, 2009) that still succeeds at twice a normal speech rate (Adanka and Devlin, 2010). To cope with this, memory traces of previous words need to be directly accessible from the current state of the language system. In this work, we propose an account of processing memory that implements these constraints—the active, online construction of meaning, sensitivity to temporal precedence, and direct access to linguistic cues. On this view, the language system is an open dynamical system that is forced by sentence inputs into transient states that correspond to their interpretations. Memory on short timescales has been conceptualized as elevated neural activity that persists beyond stimulus duration (see Durstewitz et al., 2000 for an overview). Information is encoded in spike trains and preserved in memory through sustained firing due to appropriately tuned excitatory feedback (Wang, 2001). Activity is described by the membrane voltage but there are other physiological state variables that could be harnessed for storage (Chaudhuri and Fiete, 2016). These include, for instance, synaptic currents and processes related to neuronal and synaptic adaptation. The temporal characteristics of these dynamic variables determine the lifespan of information in memory. Since these variables are physically distinct from the active membrane state, they have been termed *activity-silent* (Barak and Tsodyks, 2014; Stokes, 2015). This idea about the neurobiological basis of short-term memory is illustrated in 1. During exposure, input drives spiking activity which triggers post-synaptic potentials, adaptive effects in neuronal excitability (*intrinsic plasticity*), and short-term changes in synaptic efficacy. These effects can outlast action potentials and membrane leakage by several orders of magnitude and carry information forward on timescales that span the duration of words, phrases and whole sentences. Thus, neural spiking is viewed as an encoding operation where information is written into activity-silent dynamic variables without representing memoranda as such, consistent with observed stimulus-induced bursts of neuronal activity during encoding (Lunqvist et al., 2016). Variables of the silent state are coupled to the membrane which *continuously* reads past information back into the network’s active state. These cycles of encoding and retrieval form the neurobiological basis of a local read-write memory. The fast-changing membrane dynamics is interpreted as neural computation where numerical input is transformed to binary output, and slow-changing adaptive neural and synaptic processes serve information storage. Thus, the distinction between memory and computation can be made based on timescales. The viability of an activity-silent memory has been demonstrated by Mongillo et al. (2008) where short-term synaptic plasticity induced patterns of stimulus-specific functional connectivity from which memoranda could be retrieved after delay. The number of memorized stimuli, however, was small and recall required explicit cues. It remains open whether this approach could support real-time language processing. Here, we investigate whether processing memory could instead be provided by short-lived neuronal adaptation (Marder et al., 1996; Zhang and Linden, 2003) which has been argued to play an important role in memory formation and maintenance (Titley et al., 2017). Since language is a neurobiological system, we address this question through the simulation of neurobiologically motivated networks engaged in sentence comprehension. This *causal modeling* approach, which takes the biophysical substrate of cognitive behavior as a starting point (Koch, 1999), differs fundamentally from existing artificial neural network models of language processing (Elman, 1990; McRae et al., 1998; Ueno et al., 2011; Hinaut and Dominey, 2013; Rabovsky et al., 2018). These accounts are not modeling spike generation, neuronal adaptation, or the kinetics of synaptic transmission. Hence, they do not simulate the primitives of neural computation (e.g., action potentials and their timing) and are unable to express processing memory as history dependent changes in neuronal excitability. Crucially, these comprehension models have mainly focused on the reproduction of human behavioral data, and data fitting is often achieved through powerful, supervised learning methods whose neurobiological basis is unclear (Marblestone et al., 2016). In contrast, the causal modeling approach pursued here attempts to ground language processing in basic neurobiological design principles. In other words, rather than reproduce a specific aspect of behavior, our aim is to understand the *causal role* of these very principles. Compliance with behavioral or neuroimaging data is an independent, future outcome of this approach, not a primary modeling goal. As opposed to previous models of sentence comprehension, parameters in causal models have units of measurement that need to fall within physiological bounds. This places strong constraints on the model space and curbs arbitrary choice available in cognitive modeling. Furthermore, network time corresponds to real, physical time and this allows us to map the temporal dimension of language processing in a principled manner. Modeling the system instead of data will eventually explain how language processing and memory are implemented at the cellular level.

**Figure 1.**
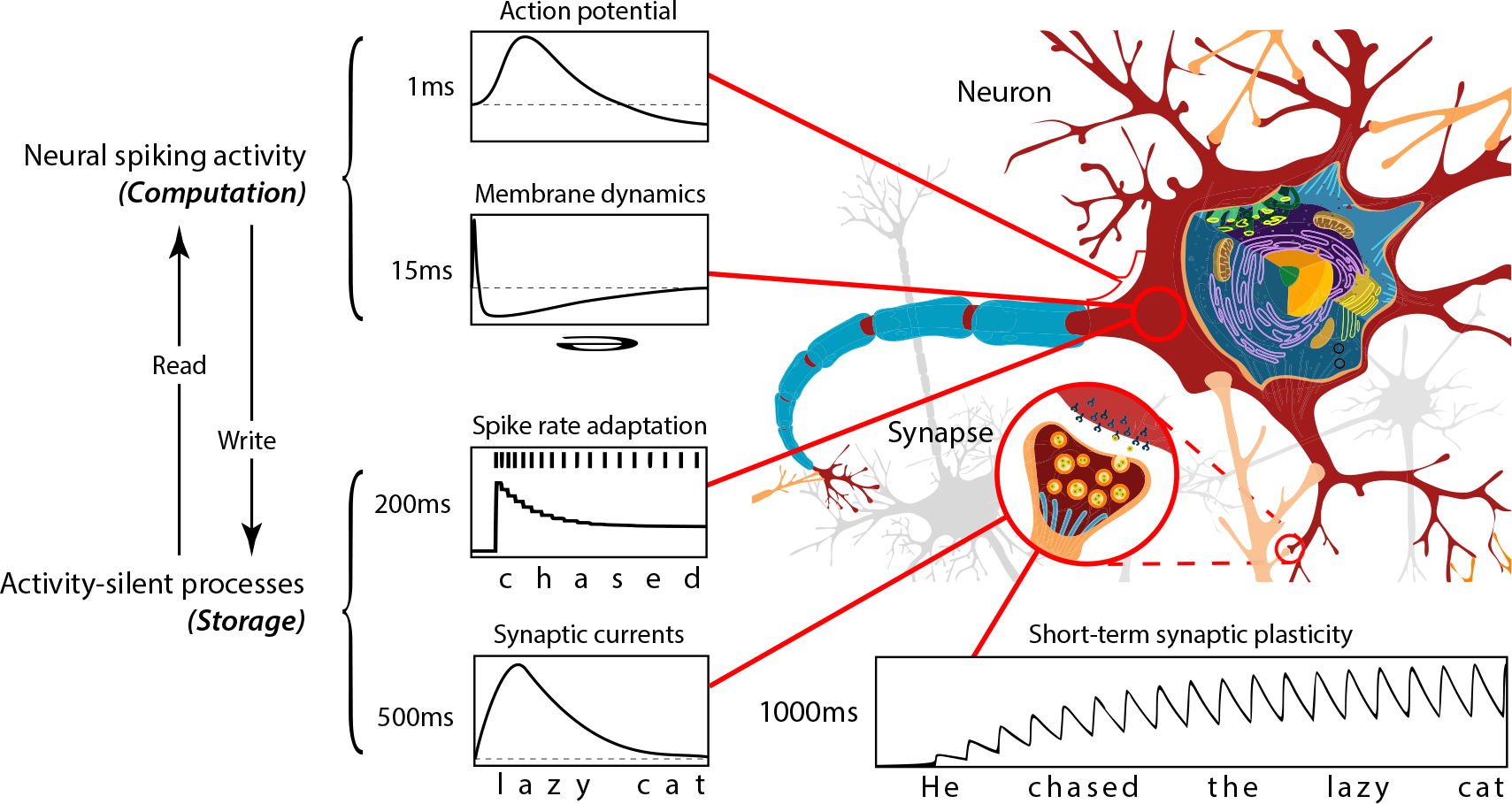
Neurobiological read-write memory. Sustained neural spiking activity has been viewed as a correlate of memory on short timescales. However, physiological processes other than the evolving membrane state can provide dynamic variables for information storage on successively longer timescales, from milliseconds to seconds. These cover the duration of basic linguistic units ranging from syllables to phrases and sentences. Coupling to neural spiking creates read-write cycles where past information, held in slow dynamic variables (storage), is continuously folded back into the fast-changing, active network state (computation). The functional segregation between memory and computation, which are co-located in neural systems, is based on the timescales of dynamic variables.

This paper is structured as follows. First, we describe a spiking network model of sentence comprehension which serves as a framework to investigate neuronal memory for language. Then, the network’s sentence processing ability and internal dynamics are characterized. Next, we demonstrate that network memory critically depends on intrinsic plasticity rather than recurrent connectivity *per se*. We examine how memory span and interference are modulated by adaptation time constants and the degree of neuronal excitability. Finally, we show that neuronal memory is suitable to establish binding relations between semantic roles and words in sentence context.

## Results

Language processing involves at least two functional components, a long-term storage of words and their phonological, morphosyntactic and semantic features which are activated by auditory or visual input (Mental Lexicon), and a combinatorial processor that integrates information into a sentence-level interpretation (Unification). Any theory of the language system needs to explain how these components are implemented and how they interact during comprehension (Hagoort, 2005; Jackendoff, 2007). Here, we focus on the modeling of the unification component which requires memory for context-dependent processing on short timescales.

### State-dependent unification network

Networks were sparsely connected circuits composed of spiking neurons with adaptation and no other plasticity (Figure 2A). 80% of neurons were excitatory and 20% inhibitory, following the ratio of pyramidal cells and interneurons in the cerebral cortex (Abeles, 1991). Information processing in these networks shares fundamental characteristics with cortical networks. First, processing results from the interaction between the input and the complete dynamic state of the network. This includes the ongoing temporal evolution of the cell membrane and ‘hidden’ aspects of network states, including dynamic processes related to neuronal and synaptic adaptation. This framework is known as *state-dependence* (Buonomano and Maass, 2009) and naturally supports context-sensitive, hierarchical and recursive processing (Petersson and Hagoort, 2012; Mante et al., 2013; Hasson et al., 2015). Secondly, word-to-word processing happens on short timescales during which neural systems do not reach stable states (Uchida and Mainen, 2003). Information must therefore be maintained in evolving neural *transients* rather than in endpoints of trajectories (e.g., attractors; Rabinovich et al., 2008). And third, transients encode information as a non-linear mixture of current input and the internal state that reflects prior processing history and is suitable for local decoding by simple readout mechanisms (Rigotti et al., 2013).

Networks for unification were exposed to a stream of time-varying language input (Figure 2A) which was generated from construction grammar templates and their syntactic alternations (e.g., *“the cat chases a toy”* versus *“a toy is chased by the cat”*; Goldberg, 2006). Synaptic projections for each input word were excitatory and random, following an exponential distribution to create heterogenous evoked dynamics. Statistically, each word targeted 5% of all neurons in the circuit and exposure times were proportional to word orthographic length. Input sequences induced a complex, spatio-temporal pattern of spiking activity. To decode the system, a logistic regression classifier was calibrated to map network states onto desired categorical output. We emphasize here that the readout units are not considered part of the unification network. They are viewed as an *external measurement device* that is used to assess the network dynamics under different neurobiological assumptions about the processing infrastructure. In the human language system, for instance, neurobiological readouts are realized by downstream recurrent networks. As a first approximation to sentence comprehension, we modeled the online, incremental assignment of thematic roles to words and phrases (e.g., AGENT, THEME, GOAL; Figure 2A). Thematic roles specify semantic relations between event participants and are part of most linguistic theories of adult meaning (Gruber, 1976; Jackendoff, 2002). Temporal bindings of words to roles can then be aggregated into a stable representation of sentence meaning. The mapping from network activity onto semantic categories provides a psycho-physical bridging relation that might allow us to explain how neurophysiological functions are linked to cognitive functions.

**Figure 2.**
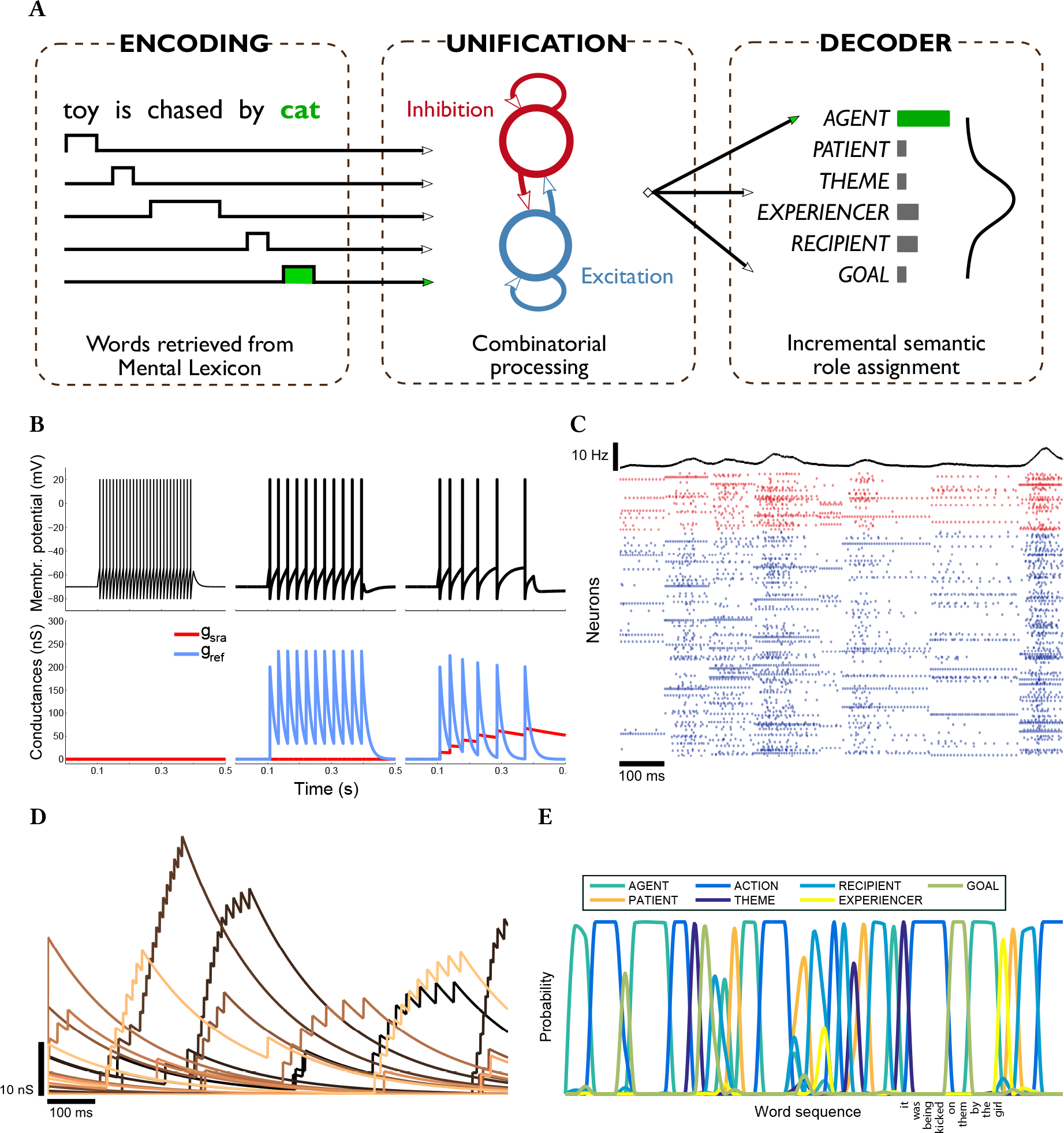
Schematic of a state-dependent unification network. **(A)** Sequences of words are mapped into a sparsely connected pool of spiking neurons with excitation and inhibition. Network states are recorded during processing and decoded onto a probability distribution over categorical semantic role labels (e.g., AGENT, THEME). In the example, the unification network needs to maintain sentence context in memory in order to assign the correct AGENT role to the sentence-final word *cat*. **(B)** Leaky integrate-and-fire neuron with spike-rate adaptation, driven by step current for a duration of 300 ms. Refractory conductance *g*_ref_ uniformly increased inter-spike intervals (middle). Spike-rate conductance *g*_sra_ adaptively decreased excitability and stretched out spike times (right). Fast, tonic spiking occurred when both conductances were turned off (left). **(C)** Raster plot and instantaneous spike rate of typical network activity at an average rate of 5 Hz. Word boundaries and neuronal adaptation are clearly visible for both excitatory (blue) and inhibitory (red) neurons as evoked responses. **(D)** Spike-triggered adaptation conductances with exponential decay recorded from the unification network during sentence processing. **(E)** Decoder semantic role response to a sequence of words. At each point in time, the most likely role for the current word is activated. Example sentence shows correct role assignment for each phrase.

Integrate-and-fire neurons in the network had a fixed voltage threshold with conductance-based mechanisms for refractoriness and spike-rate adaptation (SRA; Dayan and Abbott, 2005). SRA was modeled as a K^+^-conductance *g*_sra_ which hyperpolarized the membrane to dampen spiking activity. Following a spike, it was increased by a small amount *g*_sra_ ← *g*_sra_ + Δ*g*_sra_ and decayed back to zero exponentially with time constant *τ*_sra_ otherwise. It is the simplest linear model of spike-rate adaptation that has been described in the literature (Treves, 1993). Another conductance *g*_ref_ generated a refractory period during which neurons were prevented from spiking immediately following an action potential. Its dynamics was also modeled as an exponential decay with time constant *τ*_ref_. Both conductances modeled spike aftereffects and can be viewed as homeostatic mechanisms that counteract excessive activity by temporarily reducing excitability. They differed, however, in terms of their magnitude and time constant. While *g*_ref_ had a strong, short-term impact on the neuron, *g*_sra_ was weaker but decayed more slowly (e.g., *τ*_sra_ = 200 ms, *τ*_ref_ = 2 ms). Effects on spiking behavior exerted by these conductances are shown in Figure 2B. Whereas *g*_ref_ uniformly decreased the neuronal spike rate, *g*_sra_ adaptively slowed down spiking and the two conductances interacted via the membrane potential.

The ability to process sentences critically depends on the network’s capacity to separate distinct linguistic sequences in its transient dynamics (Maass et al., 2002). Since words were spatio-temporally coded, each input generates a distinctive pattern of evoked neural responses (Figure 2D). SRA creates history dependence in neuronal behavior as spikes trigger an instantaneous increase in the adaptive conductance which induces a current acting on the cell membrane for a duration that is proportional to the time constant *τ*_sra_. Therefore, word input can persist beyond spiking events in a neuron’s hyperpolarized membrane state. At each point in time, the degree of conductance decay provides a temporal signature for traces of contextual cues held in memory. The diversity of SRA traces in the network during sentence processing is shown in Figure 2D. Thus, the circuit maintains information as distributed, spatio-temporal representations which are characteristic of human short-term memory (Christophel et al., 2017). Memory traces of context are then used by the decoder to assign a probability distribution over semantic role labels to each input word (Figure 2E). Most target roles are activated with high confidence and competition between roles indicates early points of temporary ambiguity where sentence context does not yet fully determine semantic relations.

### Behavioral characterization and internal dynamics

To put the comprehension task into perspective, we first compared the unification network to a model with no memory of context (memoryless), and one with perfect memory (back-off n-gram). In the former, roles were regressed directly on word input. The latter model stored all n-grams from the training set, together with the target role of the n-gram final word. It also kept track of the frequency of these chunk-to-role pairs. When tested, it iteratively attempted to find the largest stored n-gram preceding the current word. Thus, the n-gram model had access to the entire sentence context in memory, and defaulted to the most frequent role in case of residual ambiguity. All models were exposed to ten random input samples of the same size and cross-validated in testing (see Methods and Materials). Memoryless regression achieved ~50% accuracy over all words but failed on the sentence final noun phrase (Figure 3A). The n-gram model reached ~75% accuracy on both measures, whereas the spiking unification network outperformed both these models, with a sentence-final accuracy of 94%. Across all words in a sentence, the network achieved ~80%. Although the n-gram model had perfect memory, with access to all-size contextual chunks and associated role frequencies in the language input, it performed less well compared to the state-dependent unification network. This indicates that the network was processing more than lexical information to map word sequences onto semantic labels. Substantial generalization from a sparse language sample points to categorical representations in the network state. This is remarkable given that input words were projected randomly into the circuit, internal connectivity was unstructured, there was no synaptic plasticity, and words shared no feature similarity with other words. This suggests that a substrate of biologically realistic neurons creates transient dynamics that can make semantic categories available to a downstream readout, in the absence of language-specific prior structure or task-related supervised learning.

**Figure 3.**
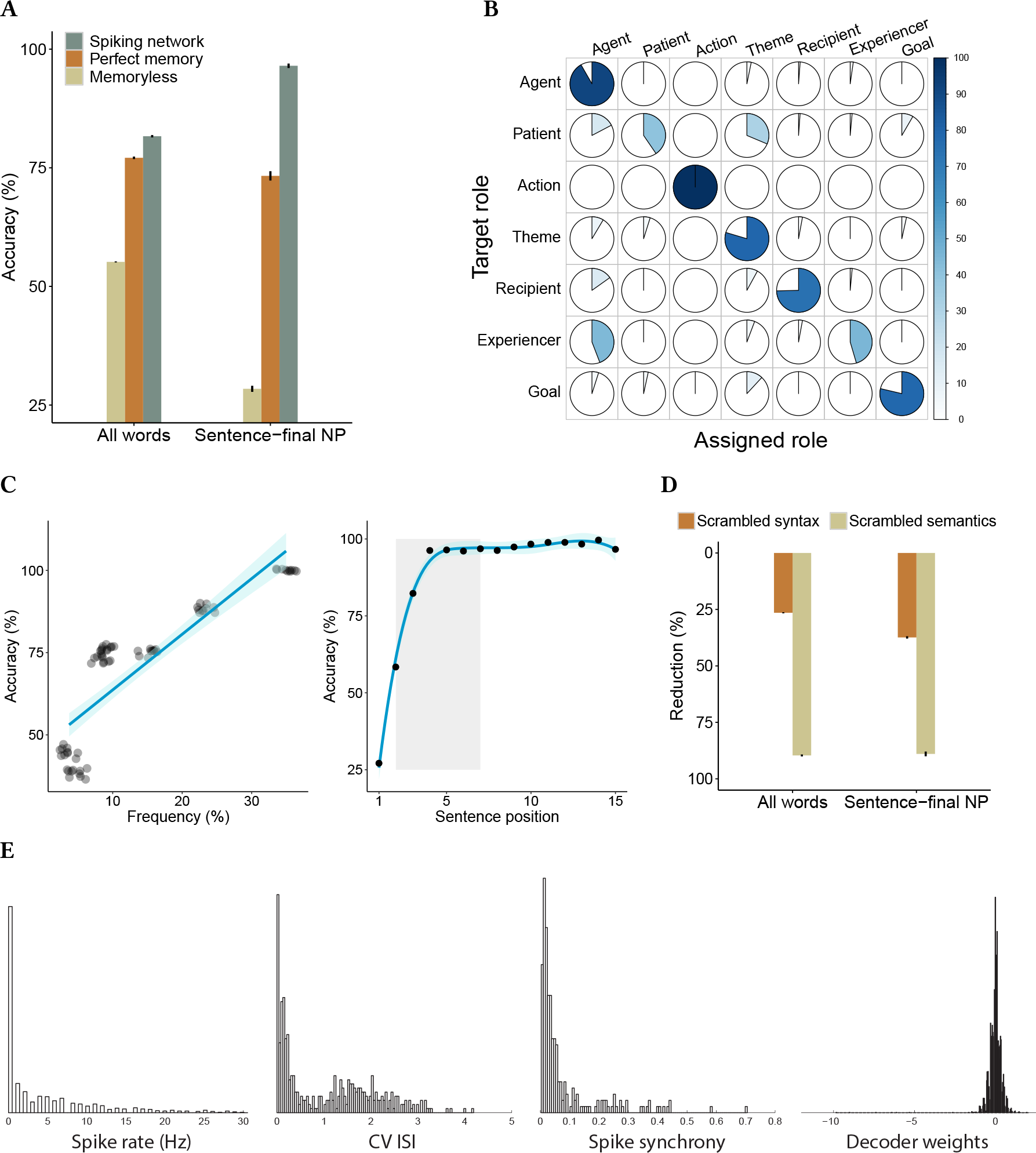
Network comprehension ability and internal dynamics. **(A)** Spiking unification network outperforms memoryless logistic regression and back-off n-gram model which has access to the entire sentence context in memory. **(B)** Network accuracy separated by semantic roles. Action labels are easy to identify, Patient and Experiencer roles are the most confusable ones. **(C)** Role assignment accuracy is highly correlated with role frequency in the input environment. Accuracy is also increasing with sentence position as contextual information is accumulating over time. Shaded area indicates verb region. **(D)** When syntax is scrambled within sentences, network accuracy reduces by 25%. When semantic roles are scrambled across sentences, accuracy reduces by 88%. **(E)** Network statistics: histograms of neuronal spike rates, coefficient of variation of inter-spike intervals (CV ISI), pair-wise spike synchrony, and distribution of readout weights (from left to right).

When performance was examined separately for each semantic role (Figure 3B), ACTION labels were identified with perfect accuracy. This is because the mapping from auxiliaries, verbs and inflections to ACTION was unambiguous. Similarly, AGENT labels were assigned with high accuracy whereas other roles (e.g., PATIENT, RECIPIENT and EXPERIENCER) were sometimes confused with AGENTS. These findings broadly reflect frequency and positional effects (Figure 3C) which are ubiquitous in human sentence processing. Labeling accuracy was correlated with role frequency (*r* = 0.86; *t* = 14.27, *p* < 0.001) and increased by sentence position. Lower accuracy in early positions reflects initial ambiguity which decreased as contextual information accumulated towards the sentence-final noun. Thus, pre-verbal, ambiguous animate nouns in less frequent roles were often treated as AGENTS which were most frequent overall.

The maintenance of contextual information relies on processing memory and assigning roles based on context, presumably, is facilitated by syntactic regularities. To identify the contribution of syntax, the network was exposed to the same language input as before but the order of words was scrambled within sentences while the mapping of words to semantic roles was kept systematic. This resulted in ~60% accuracy on both measures, a decrease of only 26% and 37% relative to baseline (Figure 3D), respectively. Thus, the gist of an utterance can still be decoded from the internal network states if context is represented in memory. Since readout projections had a large number of parameters, it was also reasonable to ask whether one could decode just about anything from the network states. To test this, semantic roles were randomized across the language input while sentences remained syntactic. This time, accuracy dropped by ~90% on both measures (Figure 3D). Hence, it was no longer possible to decode sentence meaning from contextual representations when the relation between words and their semantic roles was non-systematic.

Population activity was characterized in terms of common statistics to ensure that the unification network operated within a realistic dynamic regime during sentence processing (Figure 3E). The firing rate distribution showed a heavy-tail which is typical of cortical neurons, with some regular firing (coefficient of variation of inter-spike intervals, CV ISI, close to 0), and input-driven, irregular bursting (CV ISI above 1). Spike synchrony across the neural population was low (see Methods and Materials for details). These observations are consistent with previous findings that SRA supports the asynchronous irregular regime (Destexhe, 2009). Evoked asynchronous irregular activity has been observed in vivo (Softky and Koch, 1993; Stiefel et al., 2013) and has been argued to play a critical role in cortical information processing (van Vreeswijk and Sompolinsky, 1996; Vogels et al., 2011). Estimated readout weights were normally distributed around mean zero, with a few large, negative values. When these weights were removed, accuracy declined ~5% overall and ~1% on sentence-final noun phrases, revealing a low degree of neuronal specialization for particular semantic roles.

### Spike-rate adaptation memory

Models of short-term memory in psycholinguistics and neuroscience have been based on activity-driven maintenance of past input where excitatory feedback loops keep information alive over time. To determine the role of feedback connections on sentence comprehension ability, we compared networks with different graph topologies, synaptic density, and levels of spiking activity. We tested three hypotheses. First, if memory is provided by feedback, networks with recurrent connectivity should outperform feed-forward networks. Secondly, denser connectivity should be beneficial in maintaining sentence context since the number and length of recurrent loops grows rapidly with the number of synapses. And third, increased network activity should better utilize available feedback connections and therefore provide enhanced memory. Thus, we expected to observe an advantage of recurrent over feed-forward graphs and an increase in performance with higher synaptic density as well as higher firing rates. Results from these comparisons are shown in Figure 3A, averaged over ten randomly initialized networks per condition. To examine whether recurrence influenced role assignment accuracy, a linear mixed effects model was applied (see Methods and Materials). There was a main effect of graph type (*β* = 0.45, *χ*^2^(1) = 127.38, *p* < 0.001), indicating that network performance was lower for the recurrent graph than the feed-forward graph. Across graphs, there was a main effect of synaptic density (*β* = −0.28, *χ*^2^(1) = 28.41, *p* < 0.001), indicating that accuracy decreased with denser connectivity. There was also a main effect of network activity (*β* = −0.37, *χ*^2^(1) = 10.86, *p* < 0.001), suggesting that higher spike rates significantly impaired memory capacity. Moreover, the difference in accuracy between recurrent and feed-forward models increased with denser connectivity (*β* = 0.11, *χ*^2^(1) = 14.29, *p* < 0.001) and with higher target spike rates (*β* = 0.17, *χ*^2^(1) = 28.46, *p* < 0.001). There was no interaction between synaptic density and network activity (*β* = −0.01, *χ*^2^(1) = 0.4, *p* = 0.53). These results indicate that recurrent connectivity was not the primary source of short-term memory in our network. First, feed-forward networks outperformed recurrent ones in all conditions and this suggests that feedback did not support the maintenance of contextual information. Recurrent connectivity provided memory on very short timescales that were irrelevant for language processing where words have an extension of several hundred milliseconds. Thus, in the absence of learning, synaptic feedback appears to contribute noise to the unification task. Secondly, performance improved with sparser connectivity and decreasing spike-rates across conditions. These findings are broadly in line with studies showing that random recurrent networks have poor memory characteristics (Ganguli et al., 2008), and that feed-forward pathways can be used for structured sequence processing (Goldman, 2009).

**Figure 3.**
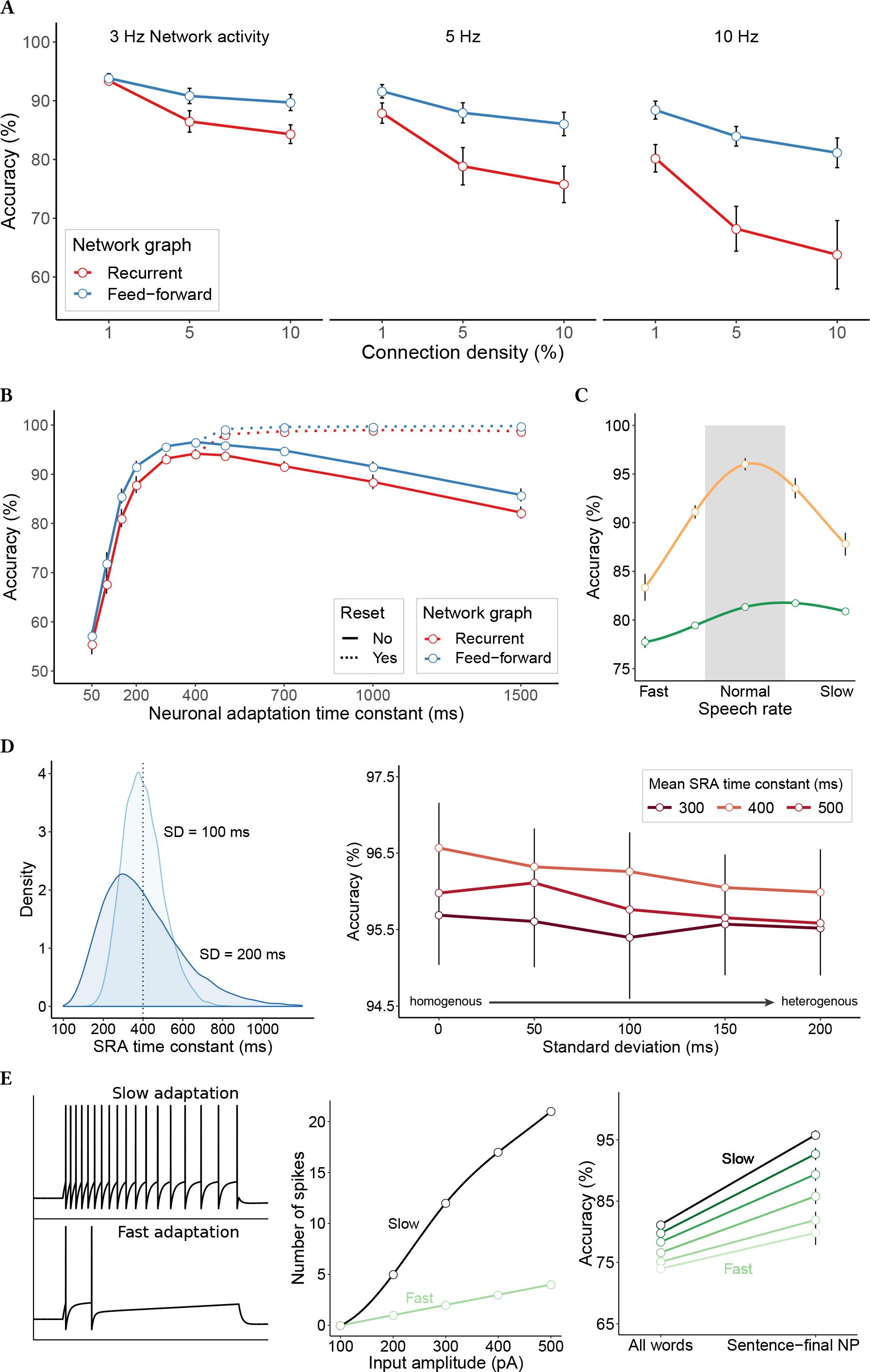
Circuit memory due to neuronal adaptation (intrinsic plasticity). **(A)** Network semantic role assignment accuracy for recurrent and feed-forward synaptic graphs with connection density varying between 1–10%, and levels of network activity between 3–10 Hz, measured on sentence-final noun phrases. Error bars show 95% confidence intervals. **(B)** Performance increases with longer spike-rate adaptation time constants and peaks for 400 ms. Reset of network dynamic variables at sentence boundaries (dashed lines) shows interference effects in processing memory. **(C)** For fixed SRA time constant, comprehension accuracy degrades when input rates are doubled (tripled) or slowed down to half (one third) of the normal speech rate. Sentence-final accuracy in orange, overall accuracy in green. **(D)** Γ-distribution of SRA time constants with mean 400 ms and standard deviations of 100 ms (light blue) and 200 ms (dark blue). Comprehension accuracy is still optimal for mean 400 ms and marginally decreases with increasing heterogeneity in adaptation time constants (right). **(E)** Fast and slow adapting neurons, controlled by the magnitude of spike-triggered increase in adaptation conductance Δ*g*_sra_ (left). Spike counts as a function of direct current input amplitude over a stimulation period of 600 ms (middle). Semantic role assignment accuracy parametrically varies with degree of neuronal excitability (right) for Δ*g*_sra_ ranging from 8 nS (slow adapting) to 500 nS (fast adapting).

The previous comparisons suggest that processing memory was provided by network features other than recurrent connectivity, with intrinsic neuronal plasticity as a candidate mechanism. To test this hypothesis, we compared networks where the time constant of conductance decay for SRA was systematically varied between 50 ms and 1.5 s. We limited these comparisons to networks with 1% connection density that were set to operate at an average instantaneous rate of 5 Hz when driven by language input (Figure 3B). Across graphs, there was an increase in comprehension ability from 50 ms to peak performance (mean: 95.4%) at 400 ms (*β* = 0.72, *χ*^2^(1) = 48.27, *p* < 0.001). Thus, the time constant of neuronal adaptation was directly related to memory span. With longer SRA time constants (above 400 ms), however, accuracy decreased significantly towards a mean of 84.0% for a conductance decay of 1.5 s (*β* = −0.32, *χ*^2^(1) = 40.63, *p* < 0.001). Longer relaxation times entail longer retention of past information and this was not always beneficial. As memory span increased, word input was eventually maintained across sentence boundaries and this adversely affected the processing of the next sentence. To demonstrate this, we reset all dynamic variables in the network at the end of each input sentence. Such rapid, cue-based clearance of the ‘hidden’ state has been observed experimentally during memory-guided behavior (Wolff et al., 2017). Figure 3B shows that accuracy continued to increase to near perfect performance (mean: 99.34%; *β* = 0.04, *χ*^2^(1) = 7.34, *p* < 0.01). Thus, flushing memory between sentences prevented traces of previous word input from contaminating the interpretation of the current sentence. This effect can be viewed as a manifestation of trace interference in short-term memory and similar contextual interference effects have been found in human language processing (Van Dyke and Johns, 2012).

The time constant *τ*_sra_ of neural adaptation is a neurobiological correlate of memory span that determines the lifetime of input traces in the network. For a time constant of 400 ms, adaptation was optimally tuned to the temporal characteristics of the comprehension task Figure 3B. This can be further validated by varying the rate of input (syllables per second) to the unification network (see Methods and Materials for details). This manipulation does not affect network dynamics but changes memory demands depending on input rate. Accuracy decreased significantly when input rates deviated from a normal rate in either direction (fast and slow), both for sentence-final words and overall comprehension ability (Figure 3C). For fast input rates, the decline was due to interference in memory, for slower rates accuracy decreased because network memory was insufficient to maintain context in longer utterances. Similar effects of speech rate variability have been found in human comprehension (Ahissar et al., 2001; Peelle and Wingfield, 2005).

Thus far, the time constant *τ*_sra_ was uniform within our circuit, although characteristics such as intrinsic plasticity might differ substantially across neuronal types and brain regions (Poulin et al., 2016; Fuzik et al., 2016). To test whether diversity in time constants affected comprehension ability, *τ*_sra_ was drawn from a Γ-distribution where the mean *μ* varied between 300–500 ms and the standard deviation *σ* between 0–200 ms (Figure 3D). These time constants were then randomly assigned to each neuron. Across *μ*, there was a decrease in performance with larger *σ* (*β* = −0.02, *χ*^2^(1) = 5.61, *p* < 0.05). When memory was cleared at sentence boundaries, however, accuracy increased (*β* = 0.34, *χ*^2^(1) = 24.71, *p* < 0.001). Hence, the decline resulted from interference due to longer timescales in the right tail of the *τ*_sra_ distribution which gradually dominated network memory with increasing *σ*. At the same time, memory loss due to the presence of shorter timescales was compensated for. This suggests that the unification network’s comprehension ability remained stable when intrinsic neuronal plasticity was heterogenous.

Apart from the timescale of intrinsic plasticity, another feature of the SRA model used here is the magnitude of spike-triggered adaptation conductances. This parameter controls how fast adaptation occurs in response to an input current. In memory parlance, it corresponds to the strength with which information is *written* into dynamic variables for storage (see Figure 1). In processing terms, when neurons are less excitable, contextual information leaves a stronger trace in the network and consequently more information is continuously read back from memory into the active network state. Figure 3E (left) shows the evolution of the membrane potential of slow and fast adapting neurons. Both are driven by the same current and have identical *τ*_sra_ but the adaptation conductance is an order of magnitude larger in the fast adapting neuron. This leads to larger spike after-hyperpolarization and a rapid decrease in excitability. The number of spikes in this neuron grows moderately as a function of input amplitude compared to the slow adapting neuron (Figure 3E, middle). Recent work has argued that transient changes in neuronal excitability, modulated by the transcription factor CREB (cAMP response element-binding), might contribute to memory function (see Lisman et al. (2018) for a review). While this proposal refers to long-term storage—the recruitment of neurons into engrams—here we tested whether changes in excitability could also have an influence on short-term processing memory. To do so, we systematically varied the magnitude of the K^+^-conductance controlling adaptation in the neuron. Since performance was dependent on spike rates (Figure 3A), network activity was kept constant by globally tuning internal connectivity strength up or down. Results show that comprehension ability was strongly modulated by the degree of neuronal excitability (*β* = −0.4, *χ*^2^(1) = 39.8, *p* < 0.001) even though adaptation time constants were fixed (Figure 3E, right). For instance, role assignment accuracy on sentence-final noun phrases dropped by 15%. These findings suggest that enhanced neuronal excitability is beneficial to processing memory for language. The precise relationship between changes in excitability, the time course of adaptation, and synaptic activity, however, remains complex and requires further analysis.

### Binding of semantic roles to words

So far, we have looked at memory characteristics that are needed to resolve thematic relations in a context-dependent manner. During online, incremental processing, the readout was binding semantic roles to words *in time*. Another important aspect of sentence comprehension is the ability to maintain these binding relations in memory until the utterance is complete. An interpretation that was adopted early in a sentence may have to be revised later on. To investigate this ability, we operationalize binding in terms of question answering (St. John and McClelland, 1990). At the end of each test sentence, the network was queried with a randomly selected semantic role label that was appropriate for this sentence. For instance, after the test item *“the toy is chased by a cat”* had been processed, the query AGENT was injected into the network (see Methods and Materials). Thus, the activity evoked by the role query was non-linearly mixed with the network’s memory of the preceding sentence. Then, a readout was estimated to map the resulting state onto the target content word for this query (Figure 4A). In the example sentence above, the correct readout response to AGENT would be *cat*. The role query acts as a semantic variable which is temporarily bound to a value by the readout, i.e., the word that fills this role slot in the test sentence. Target words could occur anywhere in the sentence and at variable distance to the position of the query.

To obtain a robust estimate of binding accuracy, networks were exposed to 20,000 sentences and tested on 5,000 queries. Since the test sentences were unique and novel, binding required substantial generalization beyond experience. Across queries, the unification network achieved ~90% role-to-word binding accuracy which is within human adult range (Wu et al., 2007). When splitting performance by queried word classes, comparable accuracy was achieved on pronouns and inanimate nouns (*β* = 0.1, *χ*^2^(1) = 1.55, *p* = 0.21), and lower accuracy on animate ones (*β* = 0.69, *χ*^2^(1) = 119.27, *p* < .001; Figure 4B). These differences are due to several factors including frequency, sentence position, degree of ambiguity, and other word distributional aspects of the input language. For instance, animate nouns could fill a larger number of distinct semantic roles than inanimate ones, and there was more variability in lexical than pronominal noun phrases. Binding accuracy was also affected by the distance between the role query (end of each sentence) and the position of the target word within the sentence (Figure 4C). Across roles, the closer the target word occurred to the query, the higher the binding accuracy (*β* = 3.65, *χ*^2^(1) = 38.5, *p* < .001). This indicates that binding was more difficult to establish the longer traces had decayed in memory. The queries most affected by distance were the AGENT, THEME, and RECIPIENT roles which all co-occurred in dative sentences. In contrast to other constructions, datives contained three noun phrases and admitted two syntactic alternations (active/passive and prepositional/ditransitive) which created positional variation and made binding relations more confusable. To trace how role-to-word bindings could be established, we inspected the network states resulting from different input queries using dimensionality reduction. This was achieved with Principal Components Analysis (PCA) on time-series of states, followed by clustering using t-Distributed Stochastic Neighbor Embedding (tSNE, see Methods and Materials). Figure 4D (left) shows that the different queries were highly separable in state space which means that the network distinctly represented the identity of the query and consequently this information was available to the readout. A similar spatial separation of semantic roles has been found experimentally in temporal cortex (Frankland and Greene, 2015). In order to answer the binding query, however, the readout also needed to uniquely identify the word that filled the corresponding role slot. When the same dimensionality reduction was performed on the states induced by a specific query (here: AGENT), different target words for this role were no longer clearly separable at the time of questioning (Figure 4D, middle). This indicates that critical information about binding relations was represented in lower variance components of the state space. To test this, readout weights were estimated on successively larger subsets of PCA space (Figure 4D, right). 76 Principle Components already explained 90% of the variance in the membrane states, corresponding to ~80% binding accuracy which still improved by ~10% when all remaining components were used in readout calibration. Thus, most of the relevant binding information was maintained in less than 10% of the dimensions of the complete dynamic state. If these findings scale to networks orders of magnitude larger, near perfect binding can be achieved by downstream readouts that are sparsely connected to subspaces of the unification network.

**Figure 4.**
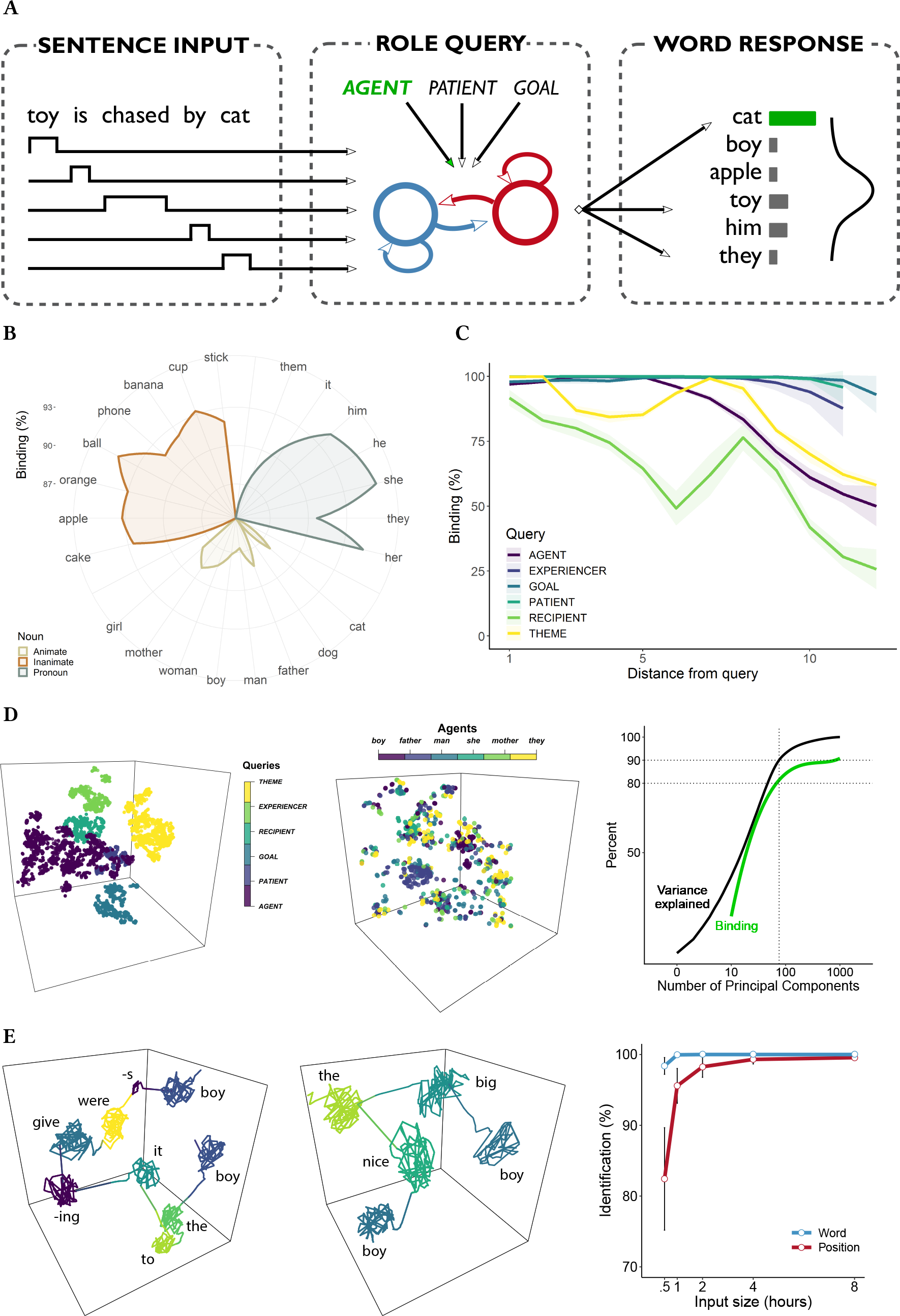
Binding of words to semantic roles. **(A)** After each input sentence, the network is queried with a semantic role label. The decoder maps the resulting network state onto a probability distribution of word responses for the queried role. A correct response occurs if the noun is identified that binds to the queried role. **(B)** Binding accuracy for different noun classes in the input language. **(C)** Binding accuracy increases with proximity to query position. **(D)** Role queries are separable in state space when combining PCA with tSNE (left). Example words that bind to the AGENT role, however, are not separable on few dimensions (middle). Variance explained versus binding accuracy for readout estimation based on PCA (right). Note the log-scale on the x-axis, dashed vertical line corresponds to 76 Principal Components. **(E)** Example sentence and its trajectory through state space (left). Multiple occurrences of the same lexical noun (*boy*) in different semantic roles (RECIPIENT, AGENT) are separated by state-dependent processing. Same noun (*boy*) preceded by different adjectives (center); separation is due to context not position. Parallel readouts decode the target word and its relative position in the sentence (right).

Binding relations are particularly challenging when utterances contain multiple occurrences of the same lexical noun in different semantic roles (‘Problem of Two’; Jackendoff, 2002). We tested this capacity in dative sentences where animate nouns could occupy both AGENT and RECIPIENT roles (e.g., *“a nice man gave the man a book”*). Inspection of sentence trajectories suggested that state-dependent processing can distinguish repeated words in memory space (Figure 4E, left). Two parallel readouts were calibrated, one that mapped role queries onto lexical fillers (as before), and another one that mapped onto the ordinal number of the word’s occurrence. Combining both readouts uniquely identifies the target noun corresponding to the queried role (e.g., in the above dative, the correct response to the query RECIPIENT would be *man–2nd*). Figure 4E (right) shows that target words and their relative position could be decoded with more than 95% accuracy in novel test items after just one hour of language input (~2,200 sentences) and eventually perfect identification was reached with longer exposure. Thus, neuronal memory can distinguish multiple occurrences of the same noun and this allows the resolution of binding ambiguities.

Establishing binding relations is a non-trivial component of sentence comprehension. It requires traces of words in memory, including a signature of temporal order (relative word position). Linguistic constructions in which these words occur (e.g., active/passive transitive) need to be distinguishable from the network state when the query is issued. This information is then combined with an identifiable semantic role key in order to deliver the correct word response. Our results indicate that all this information was present in the dynamic memory variables of the unification network. Binding relations were implicit in neuronal memory space as the network was forced by external language input into a state from which these relations could reliably be recovered. This account differs from other neural network-based approaches to binding in that it does not use specialized vector operations to form complex representations (Smolensky, 1990; Eliasmith et al., 2012), or the construction of explicit structural representations in neural tissue (van der Velde and Kamps, 2006). Moreover, our account does not require neural markers to signal binding, such as synchrony (von der Malsburg, 1995) or polychronous spiking (Izhikevich, 2006). The multi-dimensional end-state of the network’s trajectory through state space already represents the correct binding relations and this might explain why automatic language processing is so fast. In order to reason about semantic relations explicitly, downstream inference machinery can query the network and extract these relations when needed.

## Discussion

In this work we have explored the neurobiological basis of memory for language processing. We have demonstrated that neuronal adaptation can provide a context-sensitive processing memory that is suitable for online sentence interpretation and semantic binding. Contextual information was stored and maintained in activity-silent dynamic variables that regulated neuronal excitability. When intrinsic plasticity was switched to very short timescales, networks (including recurrent ones) performed poorly and this suggests that connectivity did not critically contribute to memory function. We note here that these results are not incompatible with a role of recurrence in maintenance, especially when shaped by synaptic adaptation (Markram et al., 1998) and learning (Clopath et al., 2010). Carefully balanced excitatory feedback can generate persistent activity (Amit and Brunel, 1997; Miller et al., 2003) which might serve to refresh memory traces in neuronal conductances (see Figure 1). That said, we are not aware of biophysical models for language comprehension where processing memory is provided by recurrent connectivity. Artificial network approaches (Elman, 1990; St. John and McClelland, 1990; McRae et al., 1998; Ueno et al., 2011; Hinaut and Dominey, 2013; Rabovsky et al., 2018) do not model action potentials and spike-induced forgetting, nor do they capture real-time dynamics and the temporal extension of linguistic units. If words are represented as discrete, instantaneous events, sequences are compressed in time. Each state update corresponds to one word in a sequence, whereas the real brain system evolves continuously during processing. Hence, these simplified models may be overestimating the capacity of recurrent connectivity to carry information through time. At the same time, they may be underestimating the negative effects of state variability and feedback noise due to fast membrane dynamics and spiking activity. Recurrent connectivity might play a more important role for processing than memory, when computational procedures are fine-tuned by synaptic learning. Thus, existing network models do not address the temporal mismatch between biological and behavioral timescales, and this highlights the need for a causal approach to language modeling.

### Evidence for neuronal memory

Intrinsic plasticity has been found in many types of cortical neurons (Fuhrmann et al., 2002; Koch, 1999) and has been implicated in the detection of novel input and stimulus change (Mainen and Sejnowski, 1995; Muller et al., 2007). It might serve the homeostatic regulation of network activity and prevent instability resulting from Hebbian plasticity. It has also been suggested to play a role in optimizing neuronal information transfer (Stemmler and Koch, 1999). Moreover, it might act synergistically with synaptic plasticity to support the long-term formation of memories. For instance, changes in neuronal excitability can modulate the threshold for LTP induction (Mozzachiodi and Byrne, 2010; Turrigiano, 2011; Daoudal and Debanne, 2003), and trigger consolidation. Here, we have argued that intrinsic plasticity might also play a critical role in memory on shorter timescales and we have demonstrated the functionality of such a memory system within a high-level cognitive task. This account is supported by evidence that non-synaptic, experience-dependent changes in neuronal excitability support engram formation and maintenance (Marder et al., 1996; Titley et al., 2017; Zhang and Linden, 2003; Hansel et al., 2001). Intrinsic plasticity can be expressed as a lowering of the spike-release threshold or a reduction in spike after-hyperpolarization. Both lead to higher sensitivity and an increase in firing rate. Conversely, as in SRA, excitability can decrease in response to overstimulation, causing a downregulation of output spike rates which changes the functional state of neurons. On dendritic branches, it can locally change the integration of synaptic inputs, influence coincidence detection, and selectively affect how synaptic potentials translate into action potentials (Frick and Johnston, 2005). Thus, intrinsic plasticity can temporarily alter the computational properties of neurons but also their mnemonic characteristics. Some evidence indicates that there is a direct, causal link between intrinsic plasticity and memory. Changes in excitability can function as part of the engram and serve as a transient storage device on short timescales (Titley et al., 2017; Mozzachiodi and Byrne, 2010; Zhang and Linden, 2003; Daoudal and Debanne, 2003; Marder et al., 1996). For instance, recent findings on the learning of interval durations have shown that Purkinje cells respond with temporally-specific modulations of their firing rate (over several hundred milliseconds) while circuit-internal synaptic input was pharmacologically blocked (Johansson et al., 2014). This suggests that the memory trace for temporal relations between stimuli was maintained through intracellular changes in excitability. The (de)activation of membrane conductances makes neuronal responses dependent on input history and levels of activity and short-term memory mechanisms could be based solely on such intrinsic neuronal adaptation, without changes in synaptic efficacy, or persistent activity (Marder et al., 1996).

### Memory, unification and control

The language system computes and represents complex data structures in processing memory, or unification space (Hagoort, 2005; Jackendoff, 2007). This requires the temporary binding of instances to variables in an addressable read-write memory. On the account proposed here, both reading and writing happen continuously between coupled dynamical processes that are co-located in space, but live on different timescales. Spikes in the fast membrane dynamics (*dV* / *dt* with *τ*_*m*_) trigger processes of neuronal adaptation (e.g., *dα*/*dt* with *τ*_*m*_ ≪ *τ*_*α*_), which is a *write-to-memory* operation where information is stored in slower dynamic variables *α* that are unaffected by membrane reset. The dynamic variable in which the writing occurs becomes the *physical address* of the memorandum. An initial burst of activity is sufficient to generate a memory trace that persists in the absence of sustained firing. On this account, the functional role of action potentials is to recode information into activity-silent physiological processes. Since these are coupled to the cell, memory traces continuously exert an influence on the active membrane state which corresponds to a *read-from-memory* operation (see Figure 1). These simultaneous and continuous cycles of encoding and retrieval are one correlate of neurobiological read-write memory. The rapid recoding of input into dynamic memory registers frees up computational resources for information processing. It has been proposed as a fundamental mechanism in language processing in order to deal with the fleeting nature of sensory input (Christiansen and Chater, 2015). Moreover, continuous encoding and retrieval explains the tight link between memory and the computation of meaning which happens in an online and incrementally manner within processing memory while utterances unfold in time. The degree of conductance decay endows memory traces with a *temporal signature* which is critical in sequential language processing when unification relies on word order. To control reading and writing, a functional dependence can be introduced to steer the information exchange between *V* and *α*, as in *dV* /*dt* = *f*_*V*_(*V*, *τ*_*m*_, *α*, …, 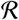) and *dα*/*dt* = *g*_*α*_(*V*, *α*, *τ*_*α*_, …, *ɛ*), where *f*_*V*_ describes the membrane evolution, *g*_*α*_ the dynamics of *α*, and *ɛ* and 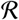 are additional dynamic variables. For instance, if *ɛ* acts multiplicatively on *V* in *g*_*α*_, *V* cannot write information into *α* when *ɛ* is near zero (e.g., via shunting inhibition; Koch, 1999). 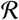 could assume a similar role in controlling retrieval. Thus, memory, unification and control in conceptual models of language (Hagoort, 2005) are mirrored in the neural microarchitecture of cognition. One aspect of language acquisition might be the learning of memory usage through control over read-write cycles. For instance, in lexicalist theories of grammar (Jackendoff, 2002) this corresponds to learning morphosyntax in the mental lexicon which specifies how words combine into larger units.

### Importance of multiple timescales

Linguistic units occupy a range of temporal grain sizes—from acoustic features to phonemes, syllables, words, phrases, clauses, sentences, and discourse—and in processing these are integrated into larger units. Since there is variation even within units, the language system needs to retain, but also discard, information on a continuum of timescales from milliseconds to seconds and minutes. These characteristics are matched by a large variety of timescales found in biological processes and this diversity may be a crucial factor in memory and information processing (Chaudhuri et al., 2014; Gjorgjieva et al., 2016).

Timescale diversity is due to different neuronal types, their sub-threshold and spike-triggered adaptive properties, the different kinetics of receptor types, and the temporal characteristics of various homeostatic and synaptic plasticity mechanisms. For instance, SRA used in our simulations is not tied to a specific time constant. Neurons can exhibit multiple adaptive traits that cover a wide range of timescales between a few milliseconds and several seconds, causing diverse neural responses (La Camera et al., 2006). This creates rich network dynamics suitable for the real-time computation of large classes of non-linear functions (Maass et al., 2007). It has also been shown that heterogeneity of responses plays an important role in context-sensitive processing which is ubiquitous in language comprehension (Rigotti et al., 2010). Furthermore, neuronal populations with different memory timescales might be recruited selectively for different task demands (Bernacchia et al., 2011). In gated recurrent networks (Cho et al., 2014), for example, neurons learn individual control functions in order to attend to different types of information in the input. Hence, they effectively operate on different timescales and this architecture copes successfully with both short and long-distance dependencies in structured sequences.

Integration establishes dependencies between units at different representational levels. A diversity of timescales might also be critical to account for such hierarchical processing. For instance, when the fast encoding dynamics of the cell membrane writes into a slow dynamic variable *α* which in turn is coupled to a slower biological process *β*, this creates a processing memory with nested timescales. If *β* implements some form of reactivation dynamics this might explain how prior information can be maintained and made available at different points in time to support hierarchical processing. Thus, processing memory for language depends on dynamic variables in physiology that move at diverse temporal scales and this heterogeneity further blurs the distinction between memory and computation. It is unlikely that this memory system is reducible to a single mechanism such as persistent activity. Future language models should incorporate multiple coupled timescales and investigate how these shape memory and hierarchical processing in sentence comprehension.

### Capacity, interference and decay

Psycholinguistic theories of memory have traditionally focused on the notion of capacity (and its limits) to explain why some sentences are more difficult to comprehend than others (e.g., Baddeley, 1986; Just and Carpenter, 1992). These accounts view memory as a passive storage space. Content is moved in and out of this buffer by a controller which mediates between memory and processor. Once the buffer is full, the resolution of dependencies begins to break down. In spiking networks, capacity would seem to be limited by available neural and synaptic resources to encode information. Our simulations suggest, however, that these were not the critical parameters that determined memory function, since it degraded with denser connectivity (increased resources; Figure 3A). Moreover, memory was highly sensitive to levels of activity and dependent on network graph (same resources). To interpret these findings, note that each word was projected into a different random subspace, generating a spatio-temporal, distributed code for the lexicon. From these input populations, activity spread into the network, differentially exciting other post-synaptic neurons. The amount of dispersion depended on neuronal out-degrees since external currents were kept constant across simulations. Stronger internal synapses and higher density entail a larger weighted out-degree. Consequently, information was diffusing into wider regions of the network and the spatial representation of words was diluted. Words became less distinguishable and this reduced the ability of the decoder to interpret the internal dynamics. Activity levels and synaptic density therefore affected the *quality* and *distinctness* of lexical representations in memory. These results support theories which argue that comprehension failure is due to interference in memory rather than limited capacity (Gordon et al., 2002; McElree et al., 2003; Lewis et al., 2006). Interference arises from representational similarity in the neural substrate (e.g., semantic interference). It is also caused when previous input is retained for too long (Figure 3B) and disrupts the interpretation of the current sentence (contextual interference). Both semantic and contextual interference have been demonstrated in human language processing (Van Dyke and Johns, 2012). And third, noise from recurrent wiring contaminated traces of lexical items in memory. These different types of interference influenced memory function, despite the fact that all networks used identical neuronal resources for storage. It has been difficult to disentangle capacity, interference and decay experimentally, and our account might help to make these concepts more precise and ground them within neurobiology.

### Conclusion

The approach presented here provides a first step towards a computational neurobiology of language. Our simulations suggest that neuronal adaptation and dynamic variables other than the membrane potentials (active state of neural systems) might be critical for our understanding of memory in language comprehension. Future work should aim to augment networks for sentence processing with models of the mental lexicon which requires plasticity mechanisms on longer timescales (Litwin-Kumar and Doiron, 2014; Zenke et al., 2015). Eventually, this causal modeling approach will bridge descriptions of brain function across levels of explanation and yield an integrated, mechanistic understanding of the human language system.

## Methods and Materials

### Neuron and synapse model

Integrate-and-fire neurons had a fixed voltage threshold with conductance-based mechanisms for refractoriness and spike-rate adaptation. The sub-threshold membrane dynamics is described by the equation

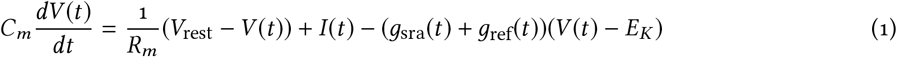

where *V*_rest_ is the resting potential, *R*_m_ denotes the leakage resistance, *C*_m_ the membrane capacitance and *I*(*t*) the total current flowing into the neuron at time *t*. When the membrane potential reached threshold *V*_th_, a spike occurred and *V* was reset to *V*_rest_. Spike-rate adaptation was modeled as a *K*^+^-conductance *g*_sra_ with reversal potential *E*_*K*_. Following a spike, this conductance was increased by a *g*_sra_ ← *g*_sra_ + Δ*g*_sra_ and it decayed back to 0 exponentially with time constant *τ*_sra_ otherwise.

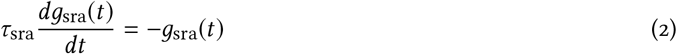

Another conductance *g*_ref_ generated a refractory period during which neurons were prevented from spiking. Its dynamics was also modeled as an exponential decay with time constant *τ*_ref_. Both conductances modeled spike aftereffects and acted homeostatically to prevent runaway activity in the network. While *g*_ref_ had a strong, short-term impact on the neuron, *g*_sra_ was weaker but decayed more slowly (e.g., *τ*_sra_ = 200 ms, *τ*_ref_ = 2 ms).

Neurons were interconnected through synapses to transmit signals. For simplicity, current-based synapses were used. The shape of synaptic currents *I*_*ij*_(*t*) was modeled as an instantaneous rise triggered by a pre-synaptic spike, followed by an exponential decay with time constant *τ*_syn_,

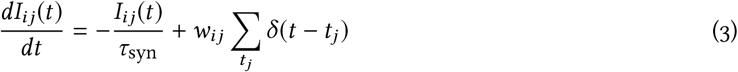

where *w*_*ij*_ is the synaptic weight from the pre-synaptic neuron *j* to the post-synaptic neuron *i*, *δ*(.) is the Dirac delta, and *t*_*j*_ are the spike times of the pre-synaptic neuron *j*. Synaptic weights *w*_*ij*_ were drawn uniformly from the real interval [0,1] ⊂ ℝ. To globally balance excitation and inhibition, inhibitory synapses were scaled to be five times stronger on average than excitatory ones. Weights were kept constant throughout the simulations, i.e., networks implemented no form of synaptic plasticity or learning. The total current into each neuron was the sum of the individual contributions of excitatory and inhibitory synaptic currents from within the network, currents generated by the adaptive conductances, and the external drive due to language input. The network was simulated with a temporal resolution of 0.2 ms and Euler’s method was used for numerical integration.

### Network graphs

Networks were composed of 1,000 neurons with 80% excitation (*E*) and 20% inhibition (*I*). Two distinct network types were compared, uniformly random recurrent graphs and feed-forward graphs where all loops were removed. Random recurrence was obtained by wiring pairs of neurons with a fixed probability that ensured a desired level of synaptic density, ranging from 1% to 5% and 10% connectivity. Thus, synapses were created independently and each one was equally likely (Gilbert random graph). Feed-forward graphs were obtained by the following procedure: a synapse *w*_*ij*_ was inserted between randomly chosen pairs of neurons *j* and *i* and the resulting directed graph was tested for cycles. If the synapse created a cycle, it was discarded, else it was retained. This procedure was iterated until the target connection density was reached. Thus, feed-forward graphs contained no cycles within *E* or *I*, nor between *E* and *I*. Experimentally matched recurrent and feed-forward networks had the same connection density and number of synapses. Network adjacency matrices were asymmetric and autapses were permitted in recurrent graphs (Bekkers, 2003).

### Language environment

As input, networks received word sequences generated from argument-structure templates with fixed patterns of function words and variation in content words (Goldberg, 2006). These *constructions* could take one, two or three verbal arguments (e.g., intransitive, transitive and dative, respectively) and included syntactic alternations when permissible. For example, the transfer dative X VERBS Y To Z could occur in prepositional or ditransitive form, in active or passive voice (e.g., *oranges are being given to a woman by the boys*). The grammar included various concepts that were organized into several categories, for example LIVING (MAN, CAT, GIRL, etc.), and OBJECT (BALL, TOY, CAKE, etc.). There were six action categories, each containing four action concepts. One of these action categories corresponded to the unergative intransitive verb class (VERB-UNERG, e.g., JUMP, DANCE), another one to the transitive class (VERB-TRANS, e.g., PUSH, HIT), and so forth. For each construction, a message template specified the set of roles that should be filled and the type of category that should fill them. The dative template, for instance, had role slots for AGENT, THEME and RECIPIENT, with AGENT and RECIPIENT roles selected from the LIVING category, and the THEME role from the OBJECT category. To create a specific message, each role was instantiated by randomly selecting a concept from the appropriate category. For example, when a message stipulated AGENT = LIVING, the AGENT role might be filled with the concept BOY. The sentence template that was paired with the message determined how that message mapped onto words and in which order. The transitive template, for example, specified that the AGENT came first, followed by an ACTION, then the PATIENT. Table 1 shows all constructions in the grammar. These sentence types and their alternations form a core subset of English. Similar to natural language, the grammar had various features such as tense (past and present), aspect (simple and progressive), and noun-verb number agreement (e.g., *“boy runs”* versus *“boys run”*). Each noun phrase was either a pronoun (e.g., *she, they, him*) or a lexical noun preceded by a determiner (optional with plurals) and an adjective (25% of the time). These features created considerable positional variation in the input sentences. Inflectional morphemes were represented as separate words (e.g., *“they were chase -ing big dog -s”*), participles and past tense were distinguished by the morphemes -*par* and -*ed*. As model input, a set of ~1500 unique sentences was generated from the grammar and concatenated into a sequence of 12,500 words. This corresponded to half an hour of language exposure in real time. Constructions were randomly selected with the same probability and instantiated from 75 lexical items in 9 word categories. Sentences were between 2 and 17 words long and the mean utterance length was 8.6 words. The construction grammar generated ~1.67 × 10^9^ distinct utterances and network input contained less than 0.0001% of the total number of sentences licensed by the grammar. Due to this expressivity of the language, substantial generalization was required from the network and processing strategies based on list memory could be excluded. Prior tests had also shown that the model was relatively insensitive to input size and the number of content words in the language, indicating robust scaling properties.

**Table 1.** Constructions in the input grammar

| Construction | Regular pattern | Syntactic alternation |
| --- | --- | --- |
| *Inanimate Intransitive* |  |  |
| Message template | ACTION = VERB-ERG; PATIENT = OBJECT |  |
| Semantic roles | PATIENT ACTION | – |
| Example sentence | *A cup was break -ing.* | – |
| *Animate Intransitive* |  |  |
| Message template | ACTION = VERB-UNERG; AGENT = LIVING |  |
| Semantic roles | AGENT ACTION | – |
| Example sentence | *Old cat -s jump -ed.* | – |
| *Transitive* |  |  |
| Message template | ACTION = VERB-TRAN; AGENT = LIVING; PATIENT = OBJECT |  |
| Semantic roles | AGENT ACTION PATIENT | PATIENT ACTION AGENT |
| Example sentence | *She kick -ss the small apple.* | *The small apple is kick -par by her.* |
| *Theme-Experiencer* |  |  |
| Message template | ACTION = VERB-THEX; THEME = OBJECT; EXPERIENCER = LIVING |  |
| Semantic roles | THEME ACTION EXPERIENCER | EXPERIENCER ACTION THEME |
| Example sentence | *They scare -ed the girl -s.* | *The girl -s were scare -par by them.* |
| *Prepositional Dative* |  |  |
| Message template | ACTION = VERB-DAT; AGENT = LIVING; THEME = OBJECT; RECIPIENT = LIVING |  |
| Semantic roles | AGENT ACTION THEME RECIPIENT | THEME ACTION RECIPIENT AGENT |
| Example sentence | *The man throw -s the stick to a dog.* | *The stick was throw -par to a dog by the man.* |
| *Ditransitive Dative* |  |  |
| Message template | ACTION = VERB-DAT; AGENT = LIVING; THEME = OBJECT; RECIPIENT = LIVING |  |
| Semantic roles | AGENT ACTION RECIPIENT THEME | RECIPIENT ACTION THEME AGENT |
| Example sentence | *The man throw -s a dog the stick.* | *A dog was throw -par the stick by the man.* |
| *Caused Motion* |  |  |
| Message template | ACTION = VERB-TRAN; AGENT = LIVING; THEME = OBJECT; GOAL = OBJECT |  |
| Semantic roles | AGENT ACTION THEME GOAL | THEME ACTION GOAL AGENT |
| Example sentence | *The boy -s were push -ing it on the cake.* | *It was push -par on the cake by the boy -s.* |
| *Locative* |  |  |
| Message template | ACTION = VERB-INTR; AGENT = LIVING; GOAL = OBJECT |  |
| Semantic roles | AGENT ACTION GOAL | GOAL ACTION AGENT |
| Example sentence | *A nice woman was drive -ing to it.* | *It was drive -par to by a nice woman.* |

### Input encoding and state collection

Words in the lexicon were encoded into random subsets of 5% of the neurons in the network such that each input word stimulated 50 neurons on average. Populations of neurons representing different words could partially overlap and, statistically, each neuron participated in the representation of 3.75 words. Thus, words were spatially coded by sparse, distributed representations but every neuron in the network was targeted by multiple words. Word input was presented to the network by injecting a step current into the target population, scaled by the corresponding input weight for each neuron. These afferent connections followed an exponential distribution with mean 0.4, and external input was assumed to be excitatory. Word exposure varied between 50 ms and 0.5 s of real, physical time, proportional to orthographic length, with a letter duration of 50 ms. When comparing speech rates (Figure 3C), this parameter varied between 18 ms and 150 ms. This corresponded to a normal speech rate of 6.1 syllables per second, a fast rate of 16.9 syllables, and a slow rate of 2.0 syllables per second. Unless stated otherwise, language input was presented as a continuous word stream without pauses or resetting of the network between sentences. While this sequence was filtered through the model, internal network states were recorded. States were defined as vectors of membrane potentials with each dimension corresponding to the current membrane voltage of one neuron. To keep simulations manageable, states were sampled at a constant rate of 200 Hz and averaged within words for each neuron. The collection of states was split into input and validation sets (10,000 and 2,500 words, respectively), and standardized before entering into a maximum entropy classifier.

### Rate tuning

Before membranes states were recorded during processing, networks were tuned to exhibit spiking activity at a target rate of 3, 5 or 10Hz within a 10% margin, respectively. First, all network-internal synapses were disconnected and synapses for external input were scaled such that networks spiked at an evoked rate of 2 Hz. Then, internal synapses were tuned to reach the overall target rate. On the first 1,000 words of the input set, the average instantaneous firing rate was determined and internal connectivity strength was iteratively adjusted up or down, accordingly, using a global synaptic scaling parameter R_app_ (Table 2).

### Decoding and evaluation

A decoder was calibrated by means of logistic regression to map network states onto target semantic roles (readout neurons). A readout is a projection of states onto the output dimensions. Typically, the projected variables are low-pass filtered spike trains. However, the filter can introduce a time constant which might distort estimation of the network’s memory characteristics (van den Broek et al., 2017). Therefore, we used filters corresponding to neuronal membrane properties and the simplest way to implement this protocol is to readout from membrane potentials. It is easily shown that this is equivalent to a real-time readout of filtered spike trains from the presynaptic neurons and hence every neuron in the network. Thus, at each point in time during online processing, reading out from membrane potentials is tantamount to reading out filtered spike trains. The decoder’s objective was to maximize the conditional data log-likelihood

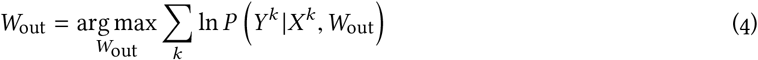

where *W*_out_ is an *M* × *N* matrix of estimated readout weights for *M* labels and *N* neurons, *X*^*k*^ is the observed network state resulting from processing the *k*th word in the training data, and *Y*^*k*^ is the target semantic role for *X*^*k*^ (Mitchell, 2017). The log-likelihood has a unique maximum but there is no closed-form solution to finding *W*_out_. To estimate these parameters, conjugate gradient descent was used with 100 iterations, and a regularization parameter of *λ* = 0.05. For discrete semantic role labels *Y* ∈ {*y*_1_, …, *y*_*M* −1_}, the probabilities *P*(*Y* = *y*_*m*_ |*X*) are then explicitly given as
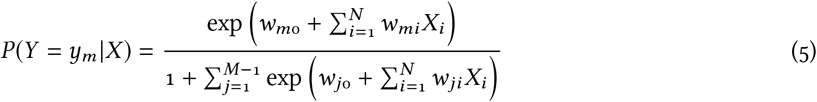

with 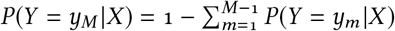. Prior comparisons had shown that logistic regression was superior to ordinary least-squares, PCA and Ridge regression.

In testing, the decoder assigned probabilities to the six semantic role labels AGENT, PATIENT, THEME, RECIPIENT, EXPERIENCER, GOAL and the ACTION label for each word in the input sentence. The role with the highest score was selected as the response and compared to the target label. Performance was quantified by means of a conservative kappa statistic *κ* for multinomial classification with *κ* = (*acc* − *rand*)/(1 − *rand*) where *acc* was the raw labeling accuracy, and *rand* the expected accuracy of a random classifier obtained through permutation of the semantic roles assigned by the decoder. Thus, the *κ* measure factors out what could be achieved by chance. We assessed role comprehension on all words in each sentence and, separately, on the sentence-final noun phrase only where memory demands on context-dependent processing were typically the highest. Model subjects were evaluated with 5-fold cross-validation on novel sets of test items. Note that the decoder itself had no memory, it could only access traces of past input that persisted in the membrane dynamics. Hence, it provided an unbiased measure of the network’s memory capacity.

### Spike regularity and synchrony

The degree of spiking regularity in the network (Figure 3E) was determined by the coefficient of variation of the inter-spike intervals (ISI)

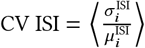

where 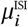 and 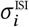 denote the mean and standard deviation of the ISIs of neuron *i* and ⟨.⟩ is the average over all neurons. The CV ISI measures spike train variability around the mean ISI. Regular spiking is indicated by a CV ISI close to 0, irregular spiking occurs for values close to 1. Values much larger than 1 indicate bursting activity. The degree of spike synchrony was quantified as the average correlation coefficient over 1,000 randomly sampled pairs of neurons

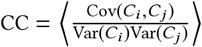

where *C*_*i*_ and *C*_*j*_ represent the number of spikes of neurons *i* and *j*, counted within successive temporal bins of 10 ms.

### Model comparison

In Figure 3A, the spiking unification network was compared to two other models, one without memory of sentence context, and one with perfect memory. In the *memoryless* model, target semantic roles were regressed directly on all the words in the training set, employing the same logistic regression classifier that was used to decode the spiking network states. The estimated model was then applied to predict the semantic roles of the words within the novel test sentences of the validation set (5 folds). The *perfect memory* model was a back-off *n*-gram learner. It was first segmenting each input sentence *S* into all possible contiguous chunks of *n* words with 1 ≤ *n* ≤ *N*, where *N* was the number of words in *S*. Each chunk was labeled with the target semantic role of the chunk-final word and these associations were stored in a hash table. Since chunks could be associated with several different roles (e.g., the word sequence *“the big cat”* could occur as an AGENT, EXPERIENCER or RECIPIENT in the training items), the learner also recorded the frequency of all chunk→role associations. When processing a test sentence *S*\* = *w*_1_*w*_2_ … *w*_*N*_ from the validation set, the model sequentially stepped through each word *w*_*k*_ of *S*\* and looked for the largest lexical chunk *w*_1_ … *w*_*k*_ in memory. If there was a match, it assigned the highest frequency role associated with this chunk to the word *w*_*k*_. If *w*_1_ … *w*_*k*_ was not matched in memory, the model would back-off to try *w*_2_ … *w*_*k*_, and so forth. In case no contextual chunk could be found in memory, the most frequent semantic role attached to the single word *w*_*k*_ was selected as the response. Thus, the *n*-gram learner had perfect memory of all sentence contexts experienced in the training environment and used a frequency-based heuristic to deal with residual ambiguity.

### Network queries

To test the binding of words to semantic roles in neuronal memory space (Figure 4A–E), the network was queried with role labels for noun phrases after every input sentence. For instance, there were two possible queries for the item *“the mouse eats the apple”*, AGENT→*mouse* and PATIENT→*apple*, and one role was randomly selected per sentence. Role queries were encoded the same way as any other lexical item in the language, with an input duration of 100 ms. The network states that resulted from processing the query were recorded for 20,000 training items, and a readout was estimated to map these states onto target words in the lexicon that filled the queried role in each of the sentences (e.g., Query = AGENT, Target = *mouse* in the example above). Then, binding accuracy was tested on 5,000 novel and unique sentences from the validation set. The neuronal time constant *τ*_sra_ was set to 1 s and memory was cleared after each sentence to minimize interference. When the same lexical noun occurred twice in a sentence (*Problem of Two*), a second, parallel readout was estimated to indicate the ordinal number of the target noun. For instance, in the test sentence *“the small boy gave a book to the big boy”*, the correct response to the query RECIPIENT would be: *boy* (lexical readout) and *2nd* (ordinal readout).

### Modeling parameters

To enable replication, we summarize network and simulation parameters in Table 2. Values refer to the model detailed in Figure 3, parametric variations (e.g., sparseness, neuronal time constants and excitability, target firing rate, input rate, etc.) are described in the main text.

**Table 2.**
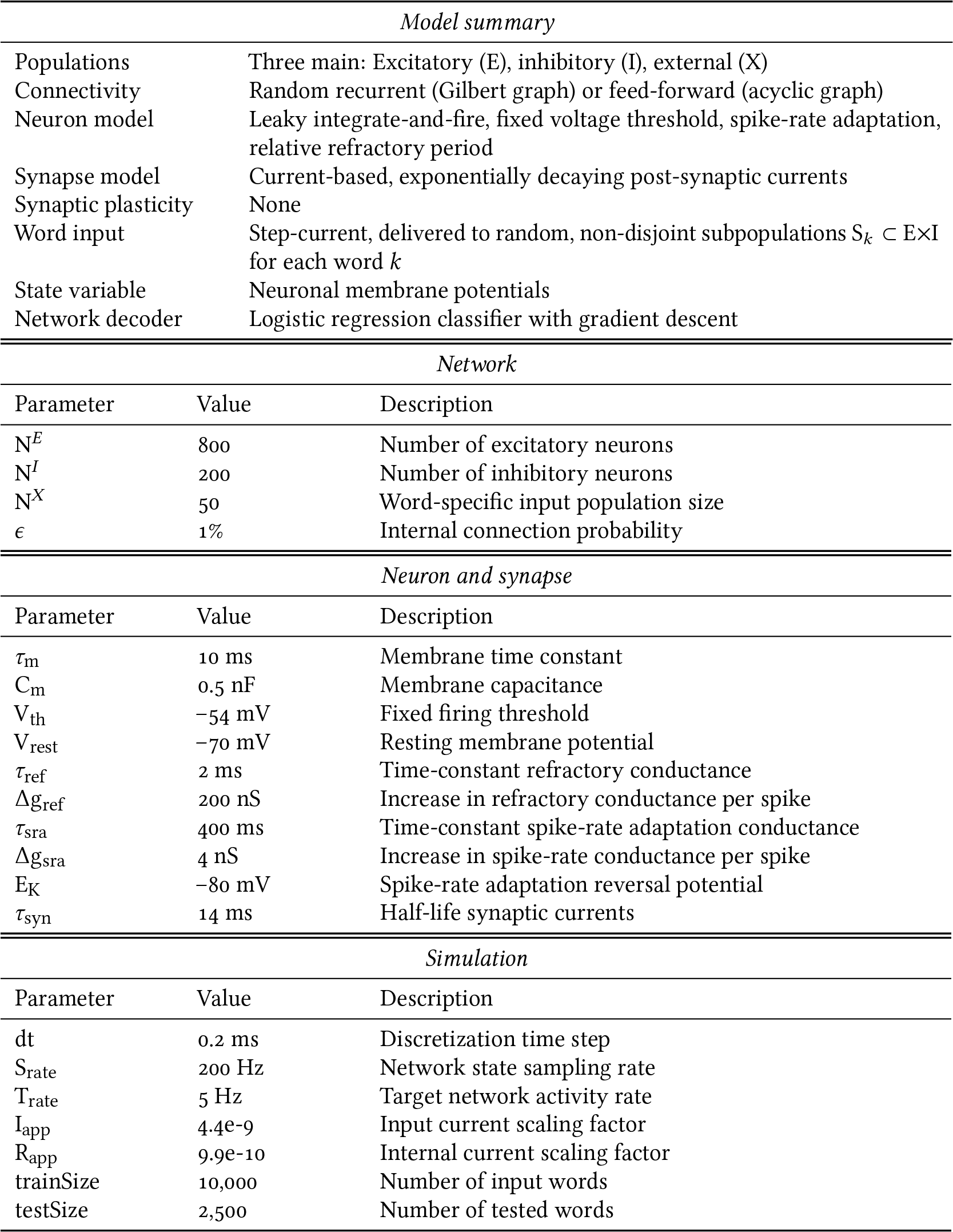
Network and simulation parameters

### Dimensionality reduction

To reduce the dimensionality of neural data for visualization (Figure 4D,E) the R implementation of tSNE (t-Distributed Stochastic Neighbor Embedding; van der Maaten and Hinton, 2008) was used on 500 initial Principal Components for 1000 iterations, with parameter values of perplexity = 30 and theta = 0.5. tSNE is a non-linear method that attempts to faithfully preserve distance between points in high-dimensional space based on a similarity metric.

### Statistical analysis

Throughout, linear mixed effects models were applied to logit-transformations of the various dependent measures described in the main text. Unless stated otherwise, categorical predictors were effect-coded, numerical ones were scaled and centered. All statistical models included the maximal random effects structure that still converged (Barr et al., 2013) and *p*-values were obtained through likelihood-ratio tests (*χ*^2^). Mixed models were run in R using the lme4 package (v. 1.1–17) with Nelder-Mead optimizer.

## Acknowledgements

This work was partially funded by the Netherlands Organisation for Scientific Research (NWO) Gravitation Grant 024.001.006 to the Language in Interaction Consortium.

## Author Contributions

Conceptualization, HF, KMP, and RD; Modeling, HF (main), MU, DvdB; Formal analysis, HF and RD; Validation, MU and DvdB; Data curation, HF, MU, and DvdB.; Writing – original draft, HF; Writing – review & editing, HF, KMP, MU, RD, and PH; Visualization, HF and MU; Funding acquisition & resources, PH; Supervision, KMP and HF.

